# Evaluating Internal Model Strength and Performance of Myoelectric Prosthesis Control Strategies

**DOI:** 10.1101/194225

**Authors:** Ahmed W. Shehata, Erik J. Scheme, Jonathon W. Sensinger

## Abstract

Ongoing developments in myoelectric prosthesis control have provided prosthesis users with an assortment of control strategies that vary in reliability and performance. Many studies have focused on improving performance by providing feedback to the user, but have overlooked the effect of this feedback on internal model development, which is key to improving long-term performance. In this work, the strength of internal models developed for two commonly used myoelectric control strategies: raw control with raw feedback (using a regression-based approach), and filtered control with filtered feedback (using a classifier-based approach), were evaluated using two psychometric measures: trial-by-trial adaptation and just-noticeable-difference. The performance of both strategies was also evaluated using a Schmidt’s style target acquisition task. Results obtained from 24 able-bodied subjects showed that although filtered control with filtered feedback had better short-term performance in path efficiency (*p* < 0.05), raw control with raw feedback resulted in stronger internal model development (*p* < 0.05), which may lead to better long-term performance. Despite inherent noise in the control signals of the regression controller, these findings suggest that rich feedback associated with regression control may be used to improve human understanding of the myoelectric control system.

## I. Introduction

Decades of advancements in myoelectric signal acquisition and processing have made myoelectric controlled prostheses a promising option for upper limb amputees [1]. Nevertheless, precise real-time decoding of movement intent from highly variable myoelectric signals and adequate methods of providing feedback to users remain a challenge [2]–[4]. Myoelectric signal variability can contribute to inconsistency in prosthesis control that results in unintended prosthesis movements [5]. Many research studies have tackled this issue by exploring feature extraction methods to obtain more useful and robust information from noisy myoelectric signals [6]. Time domain and frequency domain features are some of the most referenced of these features and are commonly used in conjunction with pattern recognition algorithms implemented in myoelectric control systems [7], [8]. The current myoelectric control systems can be broadly categorized as on/off control, proportional control, classifier-based control, and regression-based control [9].

On/off control strategies are used for binary control of a device, whereas proportional control strategies facilitate control over the speed of the prosthesis movement [10]. Although these types of control strategies are considered robust and have found clinical acceptance, the direct controllable number of degrees-of-freedom is limited by the number of usable independent control sites [11]. Classifier-based pattern recognition approaches are able to overcome this limitation, but are only capable of classifying movements sequentially [12], [13]. This drawback has more recently been overcome through 1) implementing various classifier “distributed topologies” by using singular classifiers to compare different combinations of classes and therefore enable simultaneous control [14], [15], but at the expense of accuracy and with limited control over speed and 2) use of regression-based myoelectric controllers [16], which enable simultaneous and independent speed control but at the expense of the robustness of single-class classifier-based approaches to unintentional changes in contraction patterns.

Feedback has also been shown to be of importance for robust control and in improving performance [17]. Feedback can be used for real-time regulation of control signals, as well as in the development of the user’s understanding of the system, known as their internal model [18]. Many researchers have explored the effect of feedback for real-time regulation on performance, but little work has explored its effect on internal model development (in part due to an inability to evaluate or quantify internal model strength) [19].

Quantifying this internal model enables the development of better control strategies by identifying which control strategies promote better understanding of the system and may therefore lead to better long-term performance. In a recent study [20], researchers investigated the effect of using two different myoelectric control strategies on user adaptation, which is one of the facets that can be used to estimate the strength of an internal model [21], [22]. Their results showed promising evidence that inherent feedback in myoelectric control strategies influences adaptation, which should in turn influence the user’s corresponding internal model.

In our work we sought to explicitly demonstrate that influence on a person’s internal models by measuring adaptation rate along with along with other factors, such as sensory noise and controller noise, which are necessary to calculate internal model strength. We used a recently developed psychophysical framework to assess the internal models developed using two myoelectric control strategies that differed in the feedback provided to the user during a multi degree-of-freedom (DOF) target acquisition task. The underlying signal processing for both control strategies was done using the same pattern recognition algorithm to ensure that only the feedback effect impacted the internal model strength. The first strategy implemented feedback-rich but noisy (variable) regression-based control, which we refer to here as *raw controller with raw feedback (RCRF)*. The other control strategy was analogous to a classifier, which provided reduced (discretized) feedback but more forgiving control, and is referred to here as *filtered controller with filtered feedback (FCFF)*. The psychometric test results detailed below support the hypothesis that the feedback-rich controller enables a low sensory uncertainty and strong internal model leading to a high adaptation rate. Conversely, performance test results indicate that the filtered classification-based controller yielded better short-term path efficiency and accuracy.

## II. Methods

### A. Tested Control Strategies

Many machine learning algorithms have been proposed to translate information in the myoelectric signals to either sequential or simultaneous control. Linear Discriminant Analysis (LDA) [23], Linear Regression (LR) [24], Support Vector Regression (SVR) [25], [26], and Artificial Neural Networks (ANN) [27] are some of the commonly used data-driven approaches used to identify myoelectric signal patterns for the purpose of control. SVR, in particular, is based in support vector machine (SVM) theory and can be used for either classification or regression control tasks. In both cases, SVM/SVR has been shown to yield performance superior to that of LDA/LR [6], [28], [26]. When employed as a regressor, the output is a kernel-based weighted mixture of the inputs, supporting simultaneous activation of more than one DOF at a time. This strategy is referred to here as RCRF due to the direct relationship between inputs and both control and feedback outputs. To approximate a classification output, while preserving the same decision space, the FCFF controller was achieved by gating all RCRF activations other than that of the DOF with highest level of activation (the classifier selected only the single most active DOF) (Fig. 1).

**Fig. 1.**
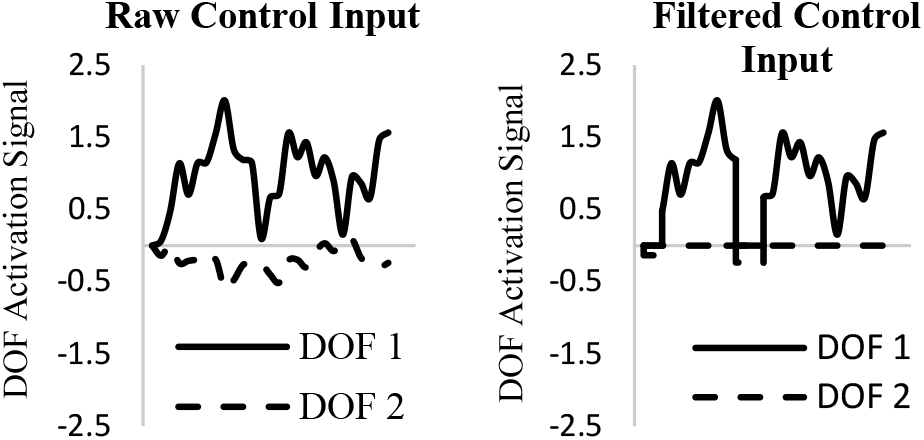
Example of control signals for 2 DOF task using Raw control and Filtered control. Raw control allows simultaneous activations of more than one DOF at a time. Filtered control selected only the single most active DOF.

### B. Experimental Setup

Subjects sat in a comfortable chair in an upright position with their line of sight perpendicular to a computer display screen. The height of the chair was adjusted to ensure a comfortable posture and that the subjects’ right arm was fully relaxed in a restrainer. This restrainer consisted of a fixed foam-padded wrist support, an adjustable foam-padded elbow support, and a foam-padded hand slot that provided resistance to hand movements while providing a comfortable setting during scheduled breaks between testing blocks. A UNB Smart Electrode System [29] was placed on the subject’s right forearm (Fig.2) and the real-time myoelectric signals extracted from muscles contractions were monitored using the Acquisition and Control Environment (ACE) software package [30] developed using MATLAB (Release 2007, The MathWorks, Inc., Natick, Massachusetts, United States).

**Fig. 2.**
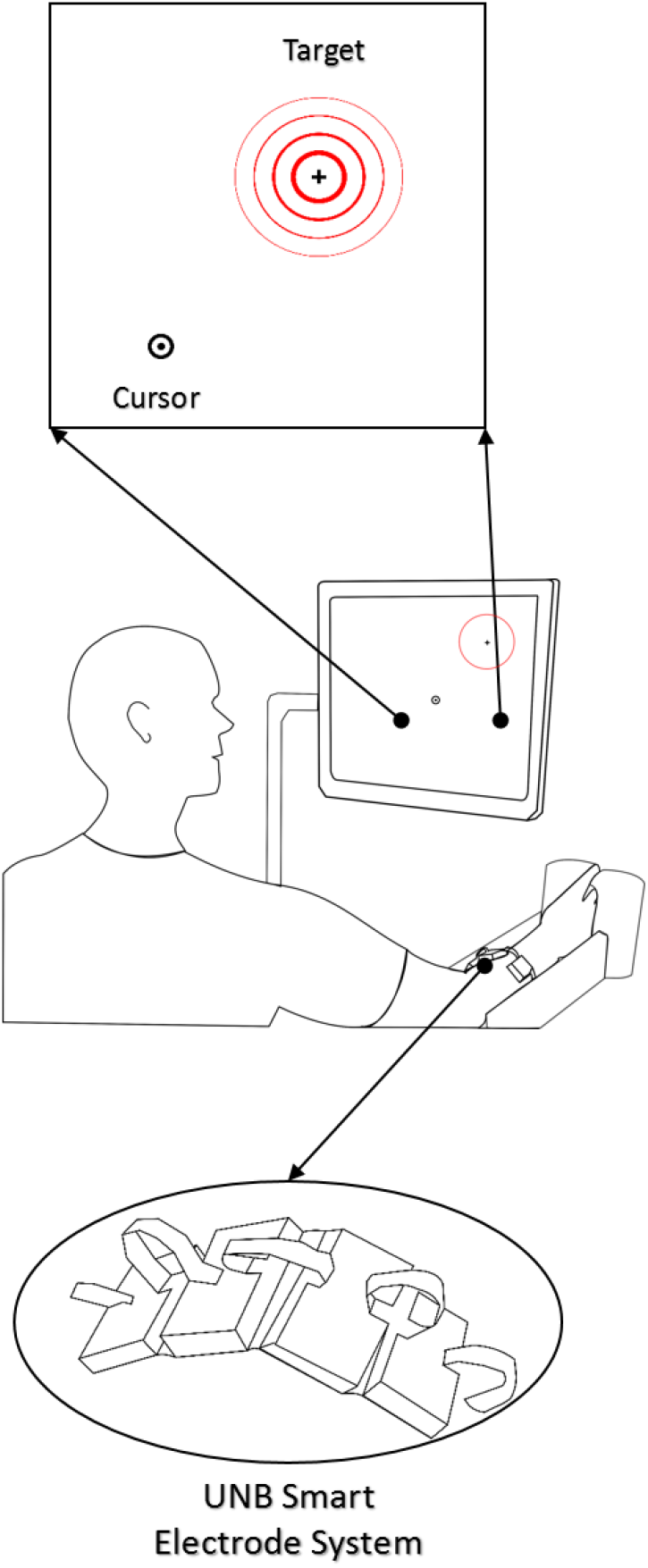
Experimental Setup. A subject performing the target familiarization block by controlling a cursor on the screen using myoelectric signals sensed by the UNB Smart Electrode System, which is placed on the subjects forearm, to acquire a target

Using isometric muscle contractions of the wrist extension/flexion and abduction/adduction, subjects controlled a cursor on the computer screen to acquire targets in a custom program that we implemented within the ACE software package. Targets consisted of cross-hairs that appeared at randomly ordered pre-determined positions on the screen. Using a Schmidt’s style test paradigm [31], a shrinking red circle of 10 pixels radius surrounded each target. This circle shrank for a pre-specified amount of time, which was determined by the testing configuration, before disappearing to indicate the end of a trial and the beginning of the next trial.

### C. Subjects

Twenty-four right-handed subjects with either normal or corrected-to-normal vision (13 male and 11 female, mean and SD of age: 25 ± 5 years) provided written consent to participate in a three-hour experimental session. These subjects were recruited primarily by word-of-mouth or the University of New Brunswick (UNB) public news and notices. Participants were informed of the overall purpose of the study, but were naive to the specific purpose and outcomes of the built-in testing blocks. The University of New Brunswick Research and Ethics Board approved this study and no compensation was provided for participation.

Eighteen of those subjects were randomly assigned to either group 1 or group 2 (nine subjects each) and the remaining subjects were assigned in a follow up study to group 3. All subjects in this study completed the same testing blocks, but the order of the control strategy presentation depended on which group a subject was assigned to. Group 1 subjects tested the RCRF control strategy first and then the FCFF control strategy, leaving group 2 subjects to start with testing the FCFF control strategy and then the RCRF control strategy.

To investigate any possible learning effect due to prolonged use of the myoelectric system, group 3 subjects acted as a control group for this study, testing the FCFF control strategy twice.

### D. Experiment Protocol

After obtaining consent from the subjects, they were asked, on a scale from 0 to 2, to rate their myoelectric control experience between no experience and moderate experience. Before placing the UNB Smart Electrode System on the subject’s right hand, the skin over their forearm was cleaned using an alcohol wipe. A couple of minutes were allowed for the real-time myoelectric signal amplitudes to settle below 1 micro-volt after which the controller model training block started. In this training, subjects were asked to follow the position of a cursor on the screen using isometric muscle contractions of the wrist for extension/flexion DOF and adduction/abduction DOF, twice for each DOF, and their myoelectric signals were recorded. An SVR pattern recognition algorithm used time-domain features extracted from these recorded signals to form the base model of the controllers used in this study. For each controller tested, subjects completed four blocks in the following order:

#### 1) Control practice and target familiarization

This block consisted of three mini tasks. For the first task, subjects were asked to ‘paint the screen’ by controlling a brush on the computer screen and trying to cover the maximum achievable area on the screen in one minute. This task was used to determine and optimize the cursor velocity mapping for each subject (20 pixels/s). Afterwards, subjects were given time to learn how to use the controller by instructing them to freely control a cursor on an empty screen starting with mild contractions in one DOF at a time and then exploring the different muscle contraction combinations and their effect on the controlled cursor for two minutes.

The last mini task consisted of three sets of 16 targets. Each target in those sets appeared at a random position from 1 of 8 predetermined positions on the screen and subjects were prompted to reach that target in 12 seconds or less. If a target was acquired in less than the indicated time, a motivational “Successful” green text appeared on the screen, the screen was cleared, and a 3 seconds count down started before another target appeared. Conversely, if the target was not acquired in the indicated time, a red “Time Out” text appeared on the screen, before beginning the countdown for the next target. Subjects were allowed to proceed to test blocks when they successfully acquired at least 75% of these targets.

#### 2) Adaptation rate test

Subjects were instructed to acquire a single target on the horizontal axis over a set of 80 trials. A trial started when the target appeared on the screen and ended after 2 seconds. Subjects were allowed 1 second to relax the contracted muscles between trials. If the target was acquired, the trial ended before the 2 seconds elapsed and provided the subject with a motivational “Successful” text, otherwise the trial terminated after the 2-second mark. Subjects were instructed to hold the cursor within a shrinking target for 200 msec for this target to be rated as successfully acquired.

#### 3) Just-noticeable-difference (JND) test

The JND is a measurement of sensory thresholds related to the estimation of specific points of the psychometric function underlying the perception of sensory stimuli [32]. In the experimental design of the JND test, the subject was forced to select between two alternative choices presented, with one of the choices having a specific stimulus added. This method is known as two-alternative forced-choice (2AFC) task, and the stimulus used was computed using an adaptive staircase [32]. This adaptive staircase quickly converges to the JND by adapting the stimulus amplitude of each trial according to the following equation:

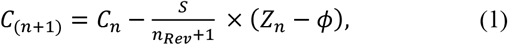

where *C* is the stimulus, *n* is the trial number, *S* is the step size, *n_Rev_* is the number of reversals between the correct and incorrect states, *Z* is a binary quantity that depends on the response at the *n_th_* trial as follows: *Z* is equal to 1 in case of success and *ø* in case of failure, and is the accuracy (set to 0.84) [33]. For this block, subjects were asked to reach a single target twice in 2 seconds and then identify which of the two trials had the added stimulus. Subjects were not given feedback on their response.

Following the (modified standard, standard) method design discussed in [34], the stimulus used was restricted to a counter-clock wise rotation of the control signal (Fig. 3). Data collected from three subjects in a pilot study was used to obtain an estimate of the initial stimulus to be used in this study. Only the target lying on the positive x-axis was explored and the test block terminated when the number of reversals reached 23 [35].

**Fig. 3.**
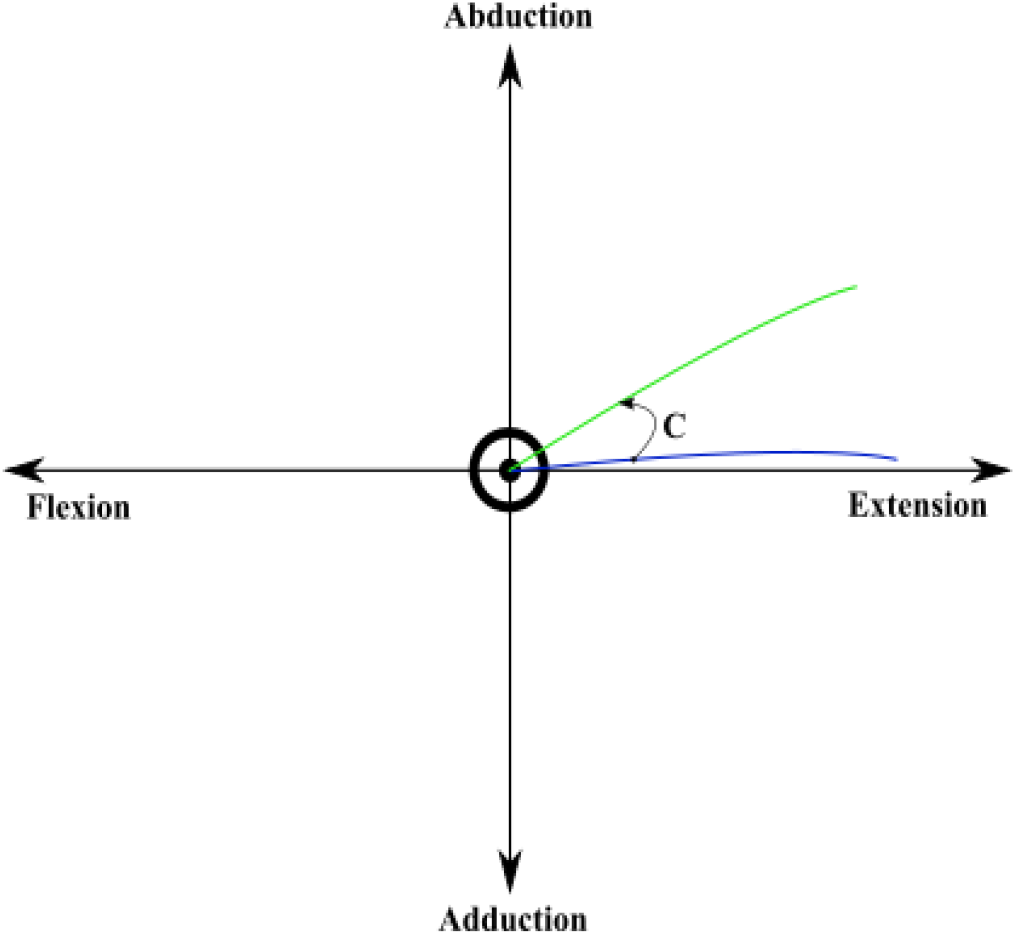
Example of a stimulus added to control signal activating cursor movement. The blue line shows actual control signal and the green line shows the perturbed control signal.

#### 4) Performance test

To assess the performance of each controller, subjects were asked to complete three challenging target sets. Each target set consisted of 16 targets which appeared at a random position drawn from 1 of 8 predetermined positions on the screen. For the first set of targets, subjects were instructed to acquire each target within 2 seconds. This time constraint was further reduced to 1.7 seconds for the second target set. The last challenge required subjects to acquire targets in 1.4 seconds or less per target. A short description of the tasks required in each block is listed in Table I.

**TABLE 1.** Summary of the test blocks

| Block | DESCRIPTION |
| --- | --- |
| Control Practice | Control a cursor on an empty screen for 2 minutes |
| Target Familiarization | Acquire 48 targets in less than 12 seconds per target<br>Targets appeared at random positions on the screen |
| Adaptation rate test | Acquire a single target on the horizontal axis in less than 2 seconds 80 times |
| JND test | Attempt to acquire a single target on the horizontal axis in less than 2 seconds twice |
| Performance test | Acquire 16 targets that appeared at random position on the screen in less than 2, 1.7, and 1.4 seconds |

### E. Outcome Measures

The main goal of this study was to assess the developed internal model strength and the performance of two commonly used myoelectric controllers. To accomplish this goal, a novel framework is used to assess internal model strength. This framework models a positioning task as a function of three variables: the sensory uncertainty (R), the control uncertainty (Q), and the internal model uncertainty (*P_param_*). For any experiment, these three variables interact to affect performance and decision, but as we show in supplementary material, their individual contributions may be extracted by collecting data for a particular set of psychophysical experiments, including 1) a trial-by-trial adaption rate test and 2) a two-alternative forced-choice test to evaluate JND. The supplementary material explains the math behind this extraction. The particular parameter of interest in this study is *P_param_*, which is a direct measure of the user’s confidence in their internal model.

#### 1) Trial-by-trial adaptation

Adaptation rate is a measure of how much the nervous system changes or modifies an internal model for a given task [36]. This rate is extracted by quantifying the rate of feedforward modification of the unfiltered control signal from one trial to the next based on error feedback from the last trial [37]. Following the same procedure used to compute the adaptation rate in [38], the following equation was used to extract this rate

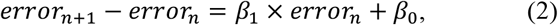

where *error* is the angle formed between the horizontal axis and the initial cursor trajectory, *n* is the trial number, *β*_0_ is the linear regression constant, and -*β*_1_ is the adaptation rate.

#### 2) JND

This parameter is a measure of the minimum perceivable stimulus in degrees identified by the subject when using each control strategy and was identified after the termination condition for the JND test has been satisfied. This parameter was used to quantify the amount of controller noise and sensory noise for each controller on a per subject basis [39].

#### 3) Internal model uncertainty *P_param_*

A novel framework was introduced to quantify uncertainties in the internal model parameters given controller noise and sensory noise parameters (see supplementary material). These uncertainties are represented in the *P_param_* parameter. The lower the value of this parameter, the higher the strength of the internal model developed.

#### 4) Performance

An experimental protocol was designed with which the short-term performance of the control strategies could be objectively evaluated using the following indicators.

##### a. Path efficiency

Paths taken to reach targets within a given time constraint during the performance test block were compared against the optimal paths [3] *i.e*., path optimizing distance covered, to compute efficiency.

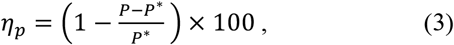

where is the path efficiency in percent, is the actual path taken, and ∗ is the computed optimal path. To ensure consistency of this measure, the optimal path for the RCRF, which enables 2-DOF simultaneous control, was computed as the shortest radial path between the cursor’s starting point and the final point where the cursor landed when the trial was terminated. For the FCFF, which allows for the activation of only one DOF at a time, the optimal path was computed as the L1 norm (Manhattan) distance (total distance travelled in X axis added to the total distance travelled in Y axis) covered to reach the final point where the cursor landed.

##### b. Accuracy

The accuracy was defined as how closely a target is reached given the time constraint. As with the technique used to ensure consistency in assessing path efficiency, the radial error was defined as the distance between the center point of a target and the actual final point reached for RCRF tests. This error was calculated as

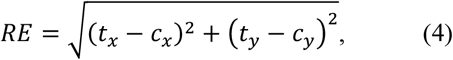

where *RE* is the radial error, *t* is the target Cartesian coordinates and *c* is the final cursor position in Cartesian coordinates. The distance error for the FCFF tests was computed as

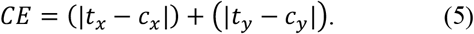

The ratio between *RE* and the shortest radial path from the starting point to the center of a target was used to compute the accuracy for the RCRF control strategy as

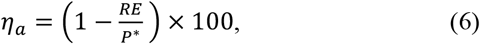

and the ratio between *CE* and the L1 norm (Manhattan) distance path from the starting point to the center of a target was used to compute the accuracy for the FCFF control strategy as

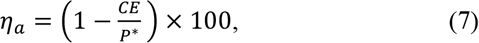

where *η*_a_ is the accuracy in percent.

### F. Data Analysis

Myoelectric signals, controller activation, and the cursor path were recorded for each trial in the test blocks. For the trial-by-trial adaptation rate, error angles were computed for the first 300–500 msec of each trial to capture the subject’s feedforward intent without incorporating feedback for real-time regulation. Only the successfully acquired target trials in the adaptation rate test were used to assess the path efficiency. Subject responses and the stimuli used in the JND test were recorded. All error bars shown in the figures were based on the standard error of the mean (SEM) to reflect its dependency on the sample size [40].

### G. Statistical Analysis

The outcome measures presented in the previous section were analyzed using two sample t-tests (equal sample size) in MATLAB and the Statistics Toolbox (Release 2014a, The MathWorks, Inc., Natick, Massachusetts, United States) to investigate the effect of controller type on each of these outcome measures. The Statistical Package for the Social Science software SPSS (IBM Corp, Released 2016, IBM SPSS Statistics for Windows, Version 24.0. Armonk, NY: IBM Corp) was used to conduct an ANOVA to investigate the effect of controller testing order on the performance and the internal model uncertainty *P_param_*. Finally, an ANOVA with repeated measures was used to compute the intraclass correlation coefficient of the adaptation rates, JNDs, internal model uncertainty, and path efficiency of group 3 subjects using a two-way mixed effects model with absolute agreement at a 95% confidence interval to investigate learning effects [41]. All analyses used a significance criterion of α = 0.05 and Leven’s test in SPSS was used to investigate homogeneity in variances of the data being analyzed to ensure that parametric test assumptions were satisfied. If they were not satisfied, nonparametric Mann-Whitney U test was used.

## III. Results

The main goal in this study was to investigate the effect of using two commonly used myoelectric control strategies on the strength of the internal model developed for a reaching task. The first control strategy, RCRF, represents a regression control strategy that allows for simultaneous control in more than 1 DOF at a time and is rich in feedback, but has comparably noisy control signals. The other control strategy, FCFF, has less variable control signals, but has reduced feedback and only allows for activation of only 1 DOF at a time. In this work the short-term performance and the strength of the internal model developed were assessed when using these control strategies. To further expand on this work, short-term performance in a target acquisition task was also evaluated when using both control strategies.

### Adaptation rate test

From equation 2, -*β*_1_ = 1 indicates perfect adaptation; lower values indicate lower adaptation; and a value higher than 1 indicates overcompensation. All subjects achieved high adaptation rates when using the feedback-rich RCRF control strategy, regardless of the order of presentation of the controllers (Fig. 4). In contrast, the order of presentation had a significant effect on the adaptation rate of subjects using FCFF after being exposed to RCRF and subjects using FCFF before being exposed to RCRF (two-sample t-test, *p* < 0.001). Upon performing t-tests on the adaptation rate data for groups 1 and 2, a significant difference was found between subjects testing RCRF in both groups 1 and 2 and subjects testing FCFF in group 2, who were not exposed to RCRF (two-sample t-test, *p* < 0.001). No significant difference was found between adaptation rate data of subjects using RCRF and subjects using FCFF after being exposed to RCRF. Exposure to the RCRF control strategy enabled subjects to adapt more when using the FCFF control strategy. This effect could be a result of either a translation of the internal model developed for RCRF to FCFF in group 1 subjects or a ramification of prolonged use of the myoelectric system, which was further investigated in more details using group 3 subjects’ results.

**Fig. 4.**
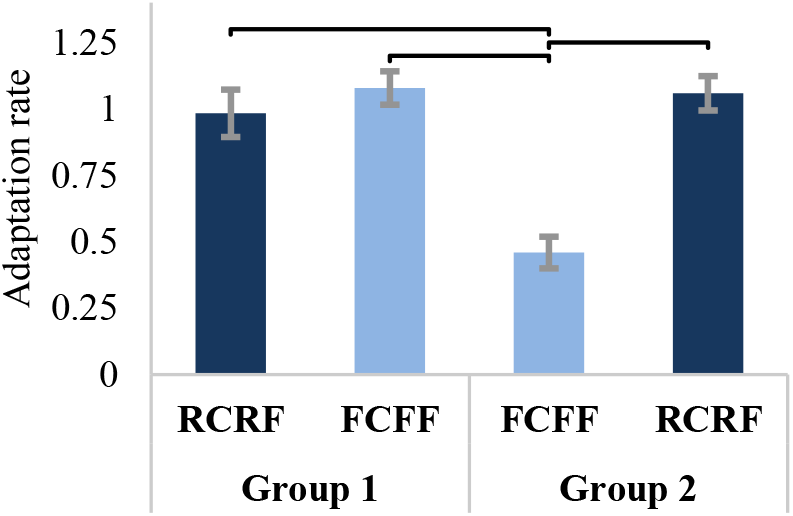
Trial-by-trial adaptation to self-generated error results across control strategies tested by subjects in groups 1 and 2. Horizontal bars indicate significant difference.

### JND test

The perception threshold for each controller was measured using the 2AFC procedure, in which an adaptive staircase was used to determine the stimulus to be added. The lower the threshold a subject was able to identify, the better ability to detect and adjust for smaller changes in the control system they had. In this test, subjects were able to identify a lower threshold when using RCRF than when using FCFF, regardless of the order of presentation (Fig. 5). Likewise, there was no significant effect due to the order of controller presentation on JND values when using FCFF. Results show that, on average, subjects were able to identify a stimulus that was at least 15∘ lower when using RCRF than when using FCFF (Fig. 6). In fact, subjects using RCRF in group 2 had a significantly lower JND value than subjects using FCFF in both groups (two-sample t-test, *p* < 0.05). Even though subjects using RCRF in group 1 obtained lower JND values than when using FCFF in the same group, this difference was not significant (two-sample t-test, *p* = 0.07). These JND values were, however, significantly lower than the values obtained by subjects using FCFF before being exposed to RCRF (two-sample t-test, *p* < 0.05). These results indicate that RCRF may enable users to identify smaller perturbations or changes in the controller by providing them with more detailed feedback.

**Fig. 5.**
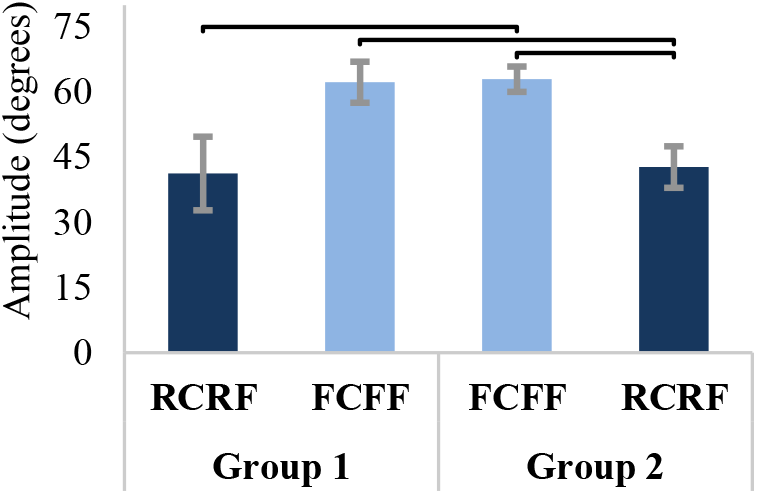
Overall JND results across control strategies for groups 1 and 2. Subjects using the RCRF control strategy were able to obtain lower JND than subjects using FCFF control strategy. Horizontal bars indicate significant difference.

**Fig. 6.**
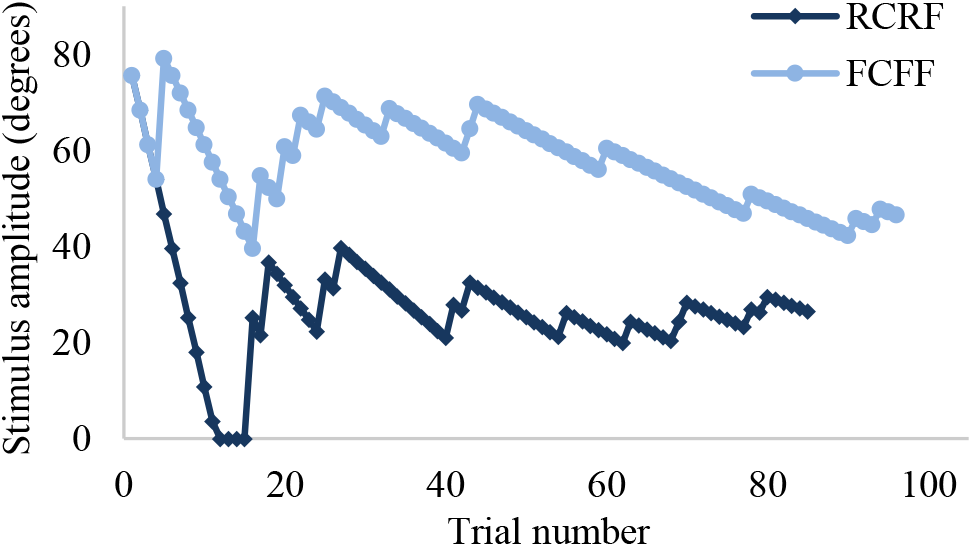
Sample data for a subject in group 1 showing the stimulus amplitude adjusted according to an adaptive staircase with a termination condition of 23 reversals and targeting 84% detection threshold.

### Internal model

Psychophysical parameters extracted from tests conducted in this study were used to quantify uncertainty in the internal model parameters developed in response to a control strategy. Quantifying this uncertainty allows for the identification of the control strategy that enables the development of a stronger internal model by indicating the one that enables lower internal model parameter uncertainty. Fig. 7 shows results for internal model uncertainty for the feedback-rich RCRF control strategy and the reduced feedback FCFF control strategy. These results show that subjects who used FCFF before being exposed to RCRF had significantly higher internal model uncertainty than subjects who used RCRF in group 1 (Mann-Whitney U test, *p* < 0.05) and RCRF in group 2 (Mann-Whitney U test, *p* < 0.01), which suggest that a feedback-rich control strategy may enable the formation of a stronger internal model.

**Fig. 7.**
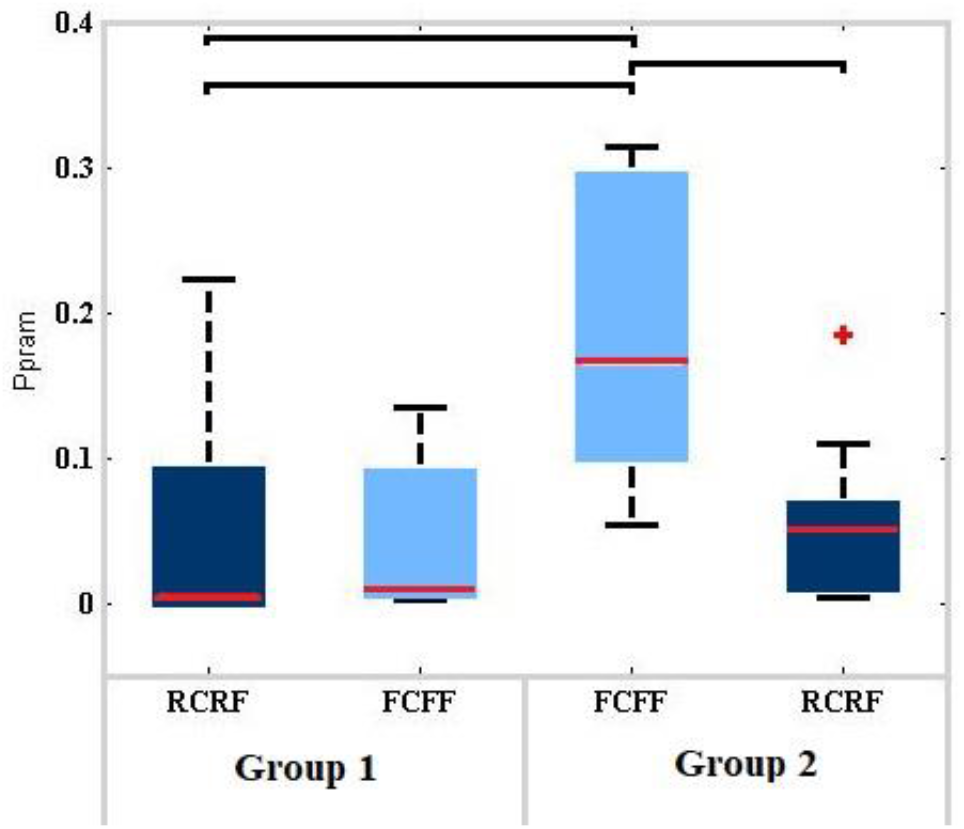
Internal model uncertainty results across control strategies for groups 1 and 2. RCRF control strategy allows for the development of a less uncertain internal model parameters than the FCFF control strategy.

In addition, subjects who used FCFF before being exposed to RCRF had significantly higher internal model uncertainty than subjects who used FCFF in group 2 (Mann-Whitney U test, *p* < 0.05). This result suggests not only that using a RCRF control strategy aids in the development of a less uncertain internal model, but also the possibility of translation of the internal model *i.e*., no significant difference between internal model uncertainty for subjects using FCFF after being exposed to RCRF and subjects using RCRF in both groups 1 and 2.

### Performance

The short-term performance of each control strategy was evaluated by determining how accurately and efficiently the task was achieved using endpoint accuracy and path efficiency. In the performance test, subjects were asked to acquire targets that were either on-axis, which optimally required the activation of 1 DOF, or off-axis, which optimally required the activation of 2 DOF simultaneously.

Results for the on-axis targets accuracy show that there was no significant difference between the control types, however the exposure to any controller had a significant effect on improving accuracy of the second controller tested. A controller that was tested in the second block outperformed the controller that was tested in the first block (two-sample t-test, *p* < 0.01) (Fig. 8. a), regardless of controller.

**Fig. 8.**
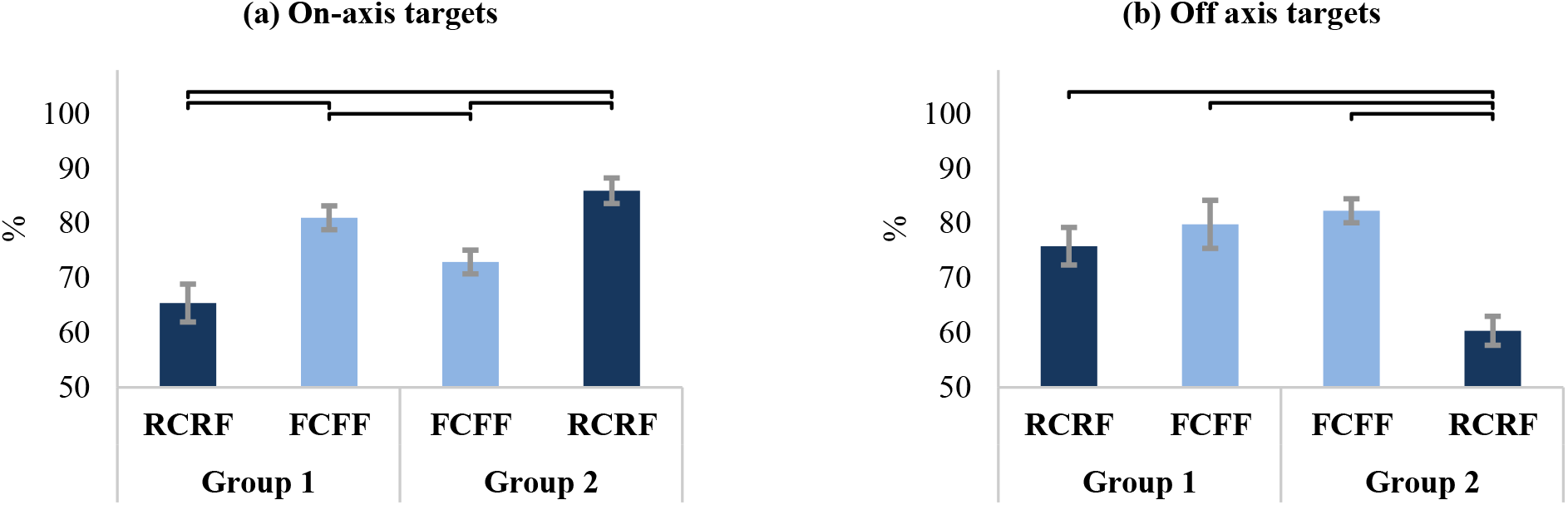
Results for accuracy normalized by the optimal distance between the starting point and the center of a target computed for each control strategy. (a) On-axis targets accuracy results show that subjects testing RCRF in group 1 achieved the lowest accuracy, but subjects in group 2 testing the same controller achieved the highest accuracy. (b) Off-axis targets accuracy results show a significant drop in the accuracy for subjects using RCRF in group 2 and all other controllers.

Interestingly, subjects using RCRF after being exposed to FCFF had significantly lower accuracy for off-axis targets than subjects using FCFF in both groups 1 and 2 (two-sample t-test, *p* < 0.01) and subjects using RCRF before being exposed to FCFF (two-sample t-test, *p* < 0.01), which suggests that subjects were influenced by the technique they used to reach off-axis targets using the sequential FCFF control strategy and therefore didn’t make use of the RCRF ability to do simultaneous movements.

The second performance assessment tool used here was path efficiency (fig. 9). Equation 3 was used to compute path efficiency for the control strategies tested in the performance test. On-axis targets path efficiency results show that for a simple 1 DOF task there was no significant difference between subjects who used RCRF first and FCFF first, however there was a significant increase in the path efficiency when retesting using any controller (two-sample t-test, *p* < 0.01). This observation may be a result of subjects exploring effective techniques that can be implemented using both control strategies. It is also worth noting the increase in the successfully acquired target count when retesting a controller. In particular, the successfully acquired target count when using RCRF after being exposed to FCFF was 5 times higher than the count for RCRF before being exposed to FCFF. This result suggests that there is a possible improvement in achieving 1 DOF tasks using the RCRF if the subject was exposed to a 1 DOF sequential control strategy.

**Fig. 9.**
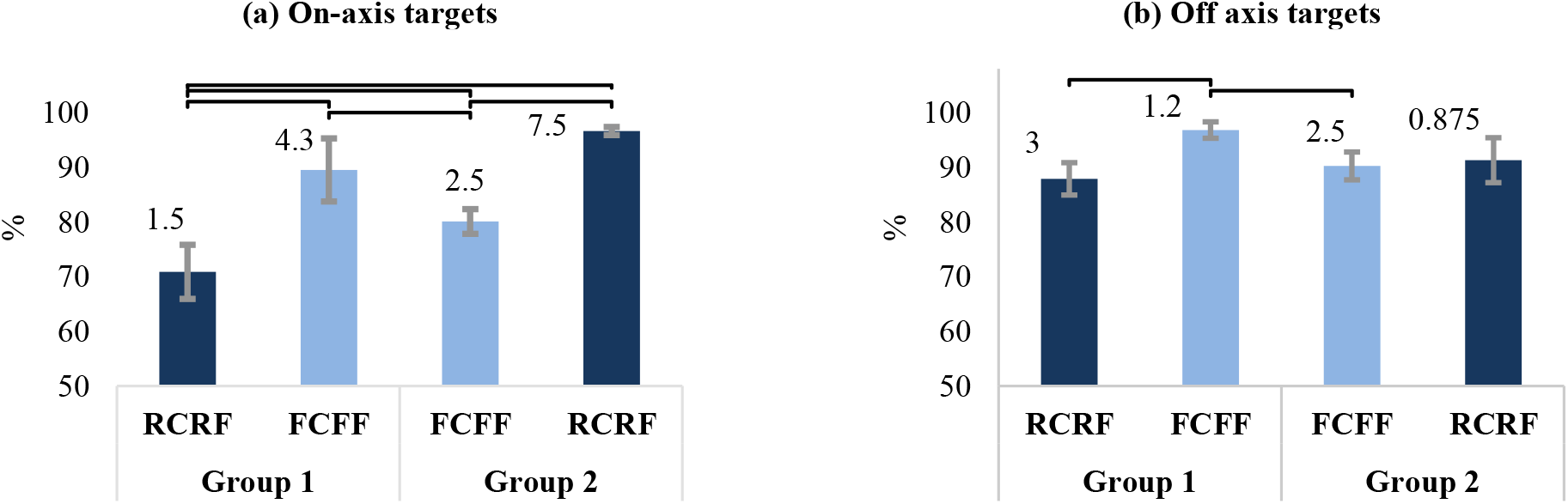
Results for path efficiency calculated with respect to the optimal path between the starting point and the final point reached computed for each control strategy. Numbers on bars show average acquired targets (maximum 8). (a) On-axis targets path efficiency results show that subjects testing RCRF in group 2 achieved the highest path efficiency and the highest success count for acquiring targets, but subjects in group 1 testing the same controller acquired the lowest success count for acquiring targets. (b) Off-axis targets path efficiency results show a significant increase in path efficiency for group 1 subjects testing FCFF after being exposed to RCRF.

For the off-axis target path efficiency, results show a significant improvement when using FCFF after being exposed to RCRF (two-sample t-test, *p* < 0.05), which may support the claim that using FCFF could be improved for a 2 DOF task if the subject was first exposed to a simultaneous control strategy like the RCRF. In addition, results show a drop in the successfully acquired target count of about 3 times when using RCRF after being exposed to FCFF.

### Group 3 results

For this group, a test-retest experiment was conducted using only the FCFF control strategy. Table II summarizes the statistical analysis for the results obtained from this group. These results show that there was no significant difference in internal model assessment parameters or performance measures when testing FCFF twice. In fact, test results for adaptation rate, JND, and internal model uncertainty showed no significant within-subject effect of retesting FCFF with good reliability (ICC > 0.6). Similarly, performance measure test results for both on-axis and off-axis targets indicated no significant difference between block 1 and block 2.

## IV. DISCUSSION

Research efforts in the field of myoelectric control have provided many solutions to improve control and performance. Classifiers, as represented here by FCFF, and regression control, as represented as RCRF, are two of the more commonly used emerging solutions. Despite differences in feedback and control signals, each of these control strategies has been found to overcome the limitations of conventional myocontrol [6], [11], [42], however their effect on the developed internal model strength, which affects user’s adaptation and the long-term performance, had not been explored. In this study, not only the short-term performance, but also the internal model strength were assessed using psychophysical and performance tests for a multi-DOF virtual target acquisition task.

**TABLE 2.** Summary of group 3 test results

| Outcome metric | ANOVA REPEATED MEASURE P | ICC | SEM |
| --- | --- | --- | --- |
| ADAPTATION rate | 0.67 | 0.74 | 0.074 |
| JND | 0.43 | 0.65 | 10 |
| Internal model uncertainty | 0.47 | 0.81 | 0.071 |
| Path efficiency On-axis targets | 0.26 | 0.72 | 1.8 |
| Path efficiency Off-axis targets | 0.79 | 0.85 | 1.9 |
| Accuracy On-axis targets | 0.81 | 0.64 | 3.2 |
| Accuracy Off-axis targets | 0.44 | 0.74 | 5.7 |

Psychophysical test results showed that RCRF enabled significantly higher adaptation to self-generated errors and the achievement of significantly lower perception threshold than FCFF. These parameters were reflected in the internal model strength, where it was also found that subjects who used RCRF developed an internal model that was significantly less uncertain (more confident) than subjects using FCFF. These results support our hypothesis that feedback-rich control strategies like RCRF enable the development of a stronger internal model than reduced-feedback control strategies like FCFF. Conversely, performance test results show that subjects had slightly better path efficiency and accuracy when using FCFF than with RCRF, which prompts a question about how much feedback is most useful for the development of a strong internal model without sacrificing the short-term performance.

The exposure to a feedback-rich control strategy enabled subjects to adapt more when using a reduced-feedback control strategy afterwards. It was found that this effect was a result of a translation of the internal model developed by the feedback-rich controller to the reduced feedback one and not a ramification of prolonged use of the myoelectric system. This phenomenon has been observed in earlier studies as structural learning [43], [44]. Test results from group 3 subjects who tested the reduced feedback control strategy twice indicated that the prolonged use of this controller did not improve performance or internal model strength.

To be able to compare performance results of the 2 DOF simultaneous RCRF control strategy with the 1 DOF sequential FCFF control strategy, the performance metrics for each control strategy were defined differently. Manhattan distances (L1 norm) were used to compute path efficiency and accuracy for the FCFF control strategy and radial (L2 norm Euclidean) distances were used to compute path efficiency and accuracy for the RCRF control strategy. This approach was used to reduce bias in the performance metrics.

Results from this study demonstrate that there was a significant improvement in the on-axis targets path efficiency and accuracy when using FCFF after being exposed to RCRF. Since the task of reaching on-axis targets only requires a simple activation of 1 DOF, the improvement in both accuracy and path efficiency may be a result of subjects using the understanding of how to effectively control the cursor in 1 DOF when they were exposed to the 2 DOF feedback-rich RCRF control strategy first and therefore informing their choice of the technique to be used to acquire targets when using FCFF. It should be noted that the exposure to any controller had a significant effect on improving accuracy of the second controller tested. From this, it may be surmised that previous experience, *i.e*., effective control technique developed and used for a control strategy, may be translated from one control strategy to another.

In contrast to our findings, Hahne et al. [20] found that classifier-based control yielded worse path efficiency than regression (they reported classifier path efficiencies of 0.27±0.12). We found similarly poor results during a preliminary investigation using LDA/LR approaches, and subsequently transitioned to SVM/SVR approaches, which have been shown to have superior performance and are equally clinically viable [16], [26]. Also in contrast to [20], which employed a position control paradigm, we used a velocity control paradigm (EMG was mapped to cursor velocity rather than position) as it is less noisy and much more commonly used in clinical practice [45], [46]. These two differences likely account for the difference in performance found between the two studies. The substantially better performance seen in this work, coupled with a more clinically relevant control paradigm likely enabled us to more accurately isolate the effects of feedback as they pertain to internal model strength.

For a grasp and lift task, researchers have investigated the effect of feedback on performance and found that feedback improved control signals [47] even after the feedback was removed [48], which was credited to the use of internal models [49]–[51]. However, these internal models were found to be unstable over time [48]. We hypothesize that this finding could be due to weakness (high uncertainty) of the internal models developed for the myoelectric controller used. Other researchers have employed psycho-physiological measurements to assess cognitive effort with and without sensory substitution methods when controlling a robot hand and found that augmenting vision with other sensory feedback reduced attentional demand [52]. We hypothesize that this reduction in attention may be due to the use of augmented feedback (feedback-rich) to build strong internal models, which are used for feedforward control and therefore enabling users to rely less on feedback for real-time regulation. The framework, psychophysical tests, and outcome measures presented here may be used to further investigate these hypotheses.

An obvious extension of this work could be using the psychophysical tests and outcome measures implemented here for a grasp-and-lift task [49] using a prosthetic hand. The assessment tools used in this work could be used in the development of new myoelectric control strategies that enable strong internal model and better performance.

In conclusion, despite classifiers such as FCFF enabling better short-term performance, regression approaches such as RCRF enabled the development of a stronger internal model, which may lead to better long-term performance [53]. With this conclusion in mind, future contributions to training and use of myoelectric prosthesis devices could be enabled by allowing users to train using feedback-rich controllers to develop a strong internal model and therefore improve long-term performance when using other controllers.

## Acknowledgements

Many thanks to Dan Blustein, Satinder Gill, Rob Smith, and Jason Robertson for thoughtful discussions. We thank Adam Wilson for continuous maintenance of the UNB Smart Electrode System.

## V. Appendix

**TABLE 3.** Summary of statistical test results for subjects using Raw control Raw Feedback in Group 1 and subjects using Filtered Control Filtered Feedback in group 2

| Outcome metric | TWO-SAMPLE T-TEST |
| --- | --- |
| ADAPTATION<br>rate | $t(16) = -4.194, p = 0.001$ |
| JND | $t(16) = -2.8, p = 0.007$ |
| Path efficiency On-axis targets | $t(16) = -2.205, p = 0.044$ |
| Path efficiency Off-axis targets | $t(16) = -0.563, p = 0.581$ |
| Accuracy On-axis targets | $t(16) = -1.68, p = 0.112$ |
| Accuracy Off-axis targets | $t(16) = -1.46, p = 0.164$ |

